# Mechanics-Driven Emergence of Mesenchymal Migration Features

**DOI:** 10.64898/2026.04.30.721940

**Authors:** Nicolas Louviaux, Ibrahim Cheddadi, Claude Verdier, Angélique Stéphanou, Arnaud Chauvière

## Abstract

Cell migration plays a central role in numerous physiological and pathological processes and emerges from the coordinated interplay between intracellular force generation, adhesion dynamics, and mechanical interactions with the environment. A minimal, mechanistically grounded understanding of these processes is required to disentangle the respective contributions of cell-intrinsic and environmental cues. Here, a two-dimensional in silico cell motility model is introduced to describe mesenchymal migration driven by intracellular traction forces generated within actin-rich protrusions anchored to a substrate. The model explicitly accounts for adhesion nucleation, maturation, force buildup and rupture, and relies on a small set of physically interpretable parameters.

A systematic mechanical analysis identifies parameter regimes that permit effective cell translocation and delineates conditions leading to stalled or mobile cells. Within motile regimes, the model reproduces a broad spectrum of cell morphologies and migratory behaviours. In particular, cell trajectories exhibit the statistical features of a persistent random walk, with a crossover from ballistic to diffusive motion that arises solely from adhesion dynamics and force balance, without imposing polarization or directional bias. Cell morphology is shown to strongly regulate migration speed, persistence, and pausing behaviour.

Altogether, this model provides a minimal reference framework for cell migration on non-deformable substrates and establishes a baseline for future studies of mechanically driven guidance. By construction, it is well suited for extension to deformable fibrous substrates, where cell-induced matrix remodeling and stiffness feedback are expected to bias migration and regulate cell encounters relevant to tissue morphogenesis and anastomosis.

## 1 Introduction

Cell migration is a complex phenomenon involved in many biological processes such as embryonic development and morphogenesis, wound healing, and cancer cell invasion. Cell migration can be triggered by biochemical growth factors, acting as chemoattractants, that define a direction to migrate towards, in the process referred to as chemotaxis. Vascular endothelial growth factor (VEGF), for example, is produced by cells experiencing hypoxia and leads to the development of new blood vessels through the migration of endothelial cells (Stéphanou et al. (2005)). Chemotaxis influences the persistence of cell motion but does not explain how cells effectively generate motion. The ability of cells to propel themselves is referred to as cell motility. In the case of mesenchymal migration, characterized by large membrane extensions, there is a broad consensus in the community on three necessary pillars enabling cell movement: cells must adhere to their environment, polarize in the direction of motion, and generate traction forces for propulsion through contraction of their cytoskeleton (DiMilla et al. (1991); Pawluchin and Galic (2022); Verdier et al. (2009); Zhu et al. (2024)).

The major component of the cellular environment is the extracellular matrix (ECM), a fibrous network mainly composed of interconnected collagen fibers (Chaudhuri et al. (2020)). Cells bind to these fibers through integrins, a family of transmembrane proteins located at focal adhesion sites. Focal adhesions are attachment protein complexes that progressively form in the peripheral region of the cell (Zaidel-Bar et al. (2004)). They are connected intracellularly to the actin network via small proteins such as talin, paxillin, and vinculin. The actin network is the main cytoskeletal component involved in mesenchymal locomotion (Svitkina (2018)). It consists of a scaffold of filaments interconnected by crosslinkers and is mostly concentrated in the peripheral region of the cell. This network results from the polymerization of actin monomers towards the cell membrane. Actin filaments interact with non-muscle myosin II, a molecular motor that induces filament sliding, leading to the formation of stress fibers that generate contraction of the actin network towards the cell nucleus. This contraction gives rise to actin retrograde flow (Small and Resch (2005)), which is restricted by focal adhesions stabilizing the actin network and allowing the cell to extend membrane protrusions such as lamellipodia. In the mesenchymal migration mode, the traction forces generated by stress fiber contraction within lamellipodia build up intracellular forces strong enough to translocate the cell nucleus; this mechanism is referred to as the molecular clutch hypothesis (Elosegui-Artola et al. (2018)). In the absence of external signals, isolated cells migrating on an isotropic substrate spontaneously exhibit random motion patterns corresponding to Lévy flights (Dickinson and Tranquillo (1993)). This type of motion is characterized by ballistic behaviour over short time intervals and diffusive behaviour over longer times. Such motion can be altered by substrate heterogeneities, such as stiffness gradients, which cells tend to follow — an experimentally observed phenomenon known as durotaxis (De Pascalis and Etienne-Manneville (2017); Shellard and Mayor (2021)). Substrate rigidity also affects cell migration speed in a biphasic manner, as measured in the study of Palecek et al. (1997) and reproduced by Scianna et al. (2013). Self-generated traction forces reach a plateau (Ghibaudo et al. (2008)) as substrate stiffness increases, with the saturation value depending on cell type (Bangasser et al. (2017)). Such substrate-dependent behaviours highlight the mechanosensing ability of cells to probe their environment and dynamically adapt their migration.

Reciprocally, cells can remodel their matrix environment by synthesizing, degrading, or deforming ECM fibers (through pulling at focal adhesions), resulting in fiber reorientation and matrix stiffening (Larsen et al. (2006)). Therefore, there exists a bidirectional interaction between cells and their environment playing a key role in migratory features and efficiency. Cell-induced ECM deformations also propagate mechanical cues that can be perceived by neighboring cells (Stéphanou et al. (2015)). Improving our understanding of the range and magnitude of this mechanical propagation would help unravel the conditions under which blood vessel sprouts successfully connect during anastomosis, a key process in vascular network growth involved in cancer development and tissue reconstruction.

To address questions related to cell migration and mechanosensitivity, many models have been proposed. Mathematical modeling is a powerful approach because models are often well-posed problems that provide unique solutions, ensuring reproducibility and allowing the use of well-established results and techniques. Most continuum models couple a mass conservation equation with reaction–diffusion–advection systems, defining quantities of interest as densities. These models can be applied across scales, from the subcellular to the tissue level (Bellomo et al. (2015)). While well suited for describing homogeneous populations following gradients (such as soluble factor concentrations), they struggle to capture one-to-one interactions, which are often treated as pointwise in such frameworks. In particular, it is difficult to incorporate the ability of cells to detect signals over a distance of multiple cell diameters (Kicheva et al. (2007)) or to represent population heterogeneity. Nevertheless, non-local models have been proposed, for instance Marchello et al. (2024), and a comprehensive review of continuum modeling approaches can be found in Chen et al. (2020). As a result, there is growing interest in agent-based models, which emphasize the representation of individuals as autonomous agents governed by interaction rules promoting emergent collective behaviours. This modeling approach allows for the description of subcellular processes driving cell biochemistry—thereby generating population heterogeneity—as well as explicit interactions between cells and their environment. Cellular Potts models (CPM) constitute a popular class of such models, describing multiscale problems through localized interaction rules. They are particularly well suited for modeling single-cell spreading and membrane deformation, with (Keijzer et al. (2025)) or without (Rens and Merks (2020)) matrix deformation, as well as collective migration during angiogenesis (Bauer et al. (2009)). However, as emphasized by the mathematical analysis of Voss-Böhme (2012), conclusions drawn from CPMs may be limited by their probability-driven nature, especially when addressing mechanical processes. Other approaches rely on phenomenological assumptions, such as the 3D model of cell motility proposed by Merino-Casallo et al. (2022), in which cells degrade the ECM to create channel-like migration paths. Similarly, Heck et al. (2020) introduced a model in which cells and ECM are represented as sets of connected particles, enabling the capture of 3D migration mechanisms within a 2D framework. In two-dimensional settings, several studies focus on ECM fiber reorientation induced by cellular traction forces (Schlüter et al. (2012); Reinhardt et al. (2013); Erhardt et al. (2025)). These modeling choices are better suited for handling spatial heterogeneities and local, dynamical interactions between system components.

However, there is always a trade-off between detailed phenomenological descriptions and the computational cost required to handle the complexity of cell motility. Consequently, other models focus on specific components of migration, such as motility initiation in one-dimensional frameworks (Wössner et al. (2024)), lamellipodia growth (Shemesh et al. (2012)), focal adhesion clustering (Liang et al. (2024)), actin cytoskeleton contraction by molecular motors (Bangasser et al. (2013)), the role of cell polarization (Dokukina and Gracheva (2010)), or a detailed representation of the ECM as a fibrous network (Mech and Rizvi (2024)). Interest in ECM modeling has grown due to the critical role of cell–ECM interactions during migration, as evidenced by the increasing number of frameworks emphasizing accurate ECM descriptions (Noël et al. (2024); Metzcar et al. (2025); Borau et al. (2024)). A comprehensive review of techniques coupling cell migration models with ECM representations can be found in Crossley et al. (2024).

A recurrent criticism of phenomenological models is the lack of emergent cell behaviours such as polarization or mechanosensitivity, which are instead imposed *a priori*. For example, the front and rear of the cell are arbitrarily defined in (Dokukina and Gracheva (2010); Heck et al. (2020)), traction forces are prescribed as functions of ECM stiffness in (Rens and Merks (2020)), or focal adhesion maturation—leading to sufficient traction forces to translocate the nucleus—is explicitly stiffness-dependent in (Shemesh et al. (2012)). The community is increasingly aware of the importance of emergence, as illustrated by the work of Bangasser et al. (2017), who proposed a molecular clutch model with emergent mechanosensing through the number of clutches and motors involved, or by Reinhardt et al. (2013), where durotaxis emerges from predefined interaction rules between cell anchors and ECM fibers.

In this work, a two-dimensional individual-based model inspired by Chauvière et al. (2024) is presented, focusing on the mechanical aspects of cell motility with minimal *a priori* assumptions. In particular, phenomenological rules such as stiffness-dependent forces or predefined cell polarization are not imposed. Instead, the model is designed to allow these properties to emerge through the probing capacity of cells when interacting with their environment. The aim of this paper is to explore thoroughly the model’s ability to reproduce various cell morphologies and dynamics. The objective is to identify key parameters enabling *in silico* cells to propel themselves and to characterize their migratory features in relation to cell morphology. The single-cell model presented here constitutes a first step towards understanding how ECM properties may subsequently be tuned to control cell protrusion dynamics and the resulting migration behaviour.

## 2 Model description

The model is inspired by Chauvière et al. (2024). It retains the general framework of the original model while providing a more accurate dynamical and mechanical description. The main components of the model are recalled below, before introducing the revised mechanical formulation in Section 2.2. A set of reference parameters is defined in Section 2.3 to non-dimensionalize the model and facilitate its analysis and calibration.

### 2.1 Model overview

#### Cell components

The cell is modeled as a set of nodes organized in a tree-like structure, rooted at the cell nucleus, with branches radiating from parent nodes. Three levels of nodes are considered, as illustrated in Fig. 1. The central node (*N*_0_) represents the cell nucleus and defines the cell position in the two-dimensional space.

**Figure 1:**
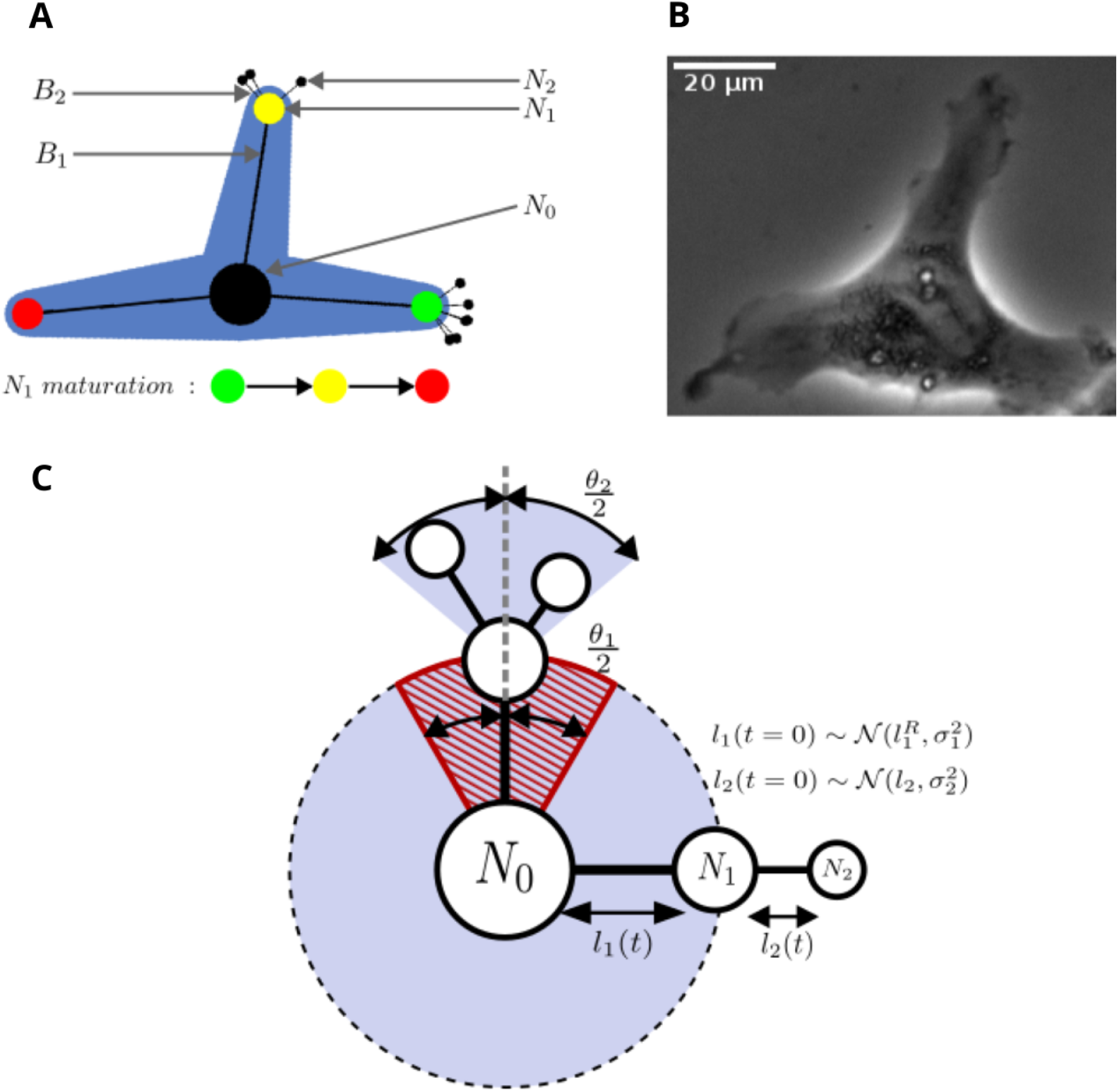
(**A**) *In silico* cell snapshot. *N*_1_ maturation process is indicated by colour: green, nascent; yellow, protrusive; red, contractile. (**B**) Phase-contrast image of an endothelial cell (EA.hy926). (**C**) Cell sketch describing cell structure control. Initial branch lengths *B*_1_ and *B*_2_ are drawn from a Gaussian distribution. The striped red area (angular width θ_1_) defines an inhibitory region for the formation of new adhesions too close to existing ones. The plain blue cone (angular width θ_2_) defines the authorized region for spawning of *N*_2_ nodes.

The second level (*N*_1_ nodes) represents adhesion complexes. *In vivo*, adhesion complexes are not localized at a single point but instead exhibit multiple anchoring points distributed in lamellipodia, each transmitting intracellular contractions to the substrate. The different contributions are assigned to a single effective force application point, the *N*_1_ node. The *N*_1_ nodes undergo a maturation process, as described in (De Pascalis and Etienne-Manneville (2017); Zaidel-Bar et al. (2004)), transitioning from nascent adhesions loosely interacting with the substrate to mature focal adhesions firmly anchored to the latter. This maturation process is decomposed into three phases: nascent, protrusive, and contractile. The connections between the nucleus and the *N*_1_ nodes model actin fibers and are represented by the *B*_1_ branches. Their mechanical properties evolve with adhesion maturation to reproduce protrusion growth and contraction (see next section). These branches transmit traction forces from adhesion complexes to the nucleus.

The third level (*N*_2_ nodes) serves a dual role of sensors and active force generators. These nodes are assimilated to filopodia tips that probe the extracellular environment and exert pulling forces on *N*_1_ nodes, enabling their displacement relative to the substrate and promoting lamellipodium formation. The *N*_2_ nodes are anchored to the substrate and have a short lifetime compared to other characteristic timescales, ensuring rapid turnover and effective probing. Each *N*_2_ node is connected to its parent *N*_1_ node by an elastic branch, denoted *B*_2_, which is initially under tension. This configuration generates an outwards force that promotes protrusion growth. The elastic behaviour of the *B*_2_ branches also endows the *N*_2_ nodes with mechanosensitive properties on deformable substrates: substrate deformation induced by pulling reduces the effective traction force, limiting protrusion growth and biasing exploration towards stiffer regions. This motivates the inclusion of *B*_2_ branches, although in the present work the substrate is assumed to be non-deformable.

#### Cell dynamics

The cell position is defined by the location of the nucleus (*N*_0_). Adhesion complexes (*N*_1_ nodes) emerge stochastically around *N*_0_ in the cell periphery. Their formation is spatially constrained by an inhibitory region surrounding existing *N*_1_ nodes (see Fig. 1C). The forces 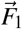 transmitted by each *B*_1_ branch to the nucleus generate a mechanical “tug-of-war” that may result in nucleus translocation. By default, the nucleus is anchored to the substrate due to static friction. When the norm of the net force exceeds a rooting threshold 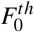, the nucleus slides with velocity 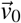, hindered by a dynamic friction coefficient modeling cell–substrate interactions during motion (Fig. 2A). The nucleus position is updated using a quasi-static approximation of the force balance acting on *N*_0_.

**Figure 2:**
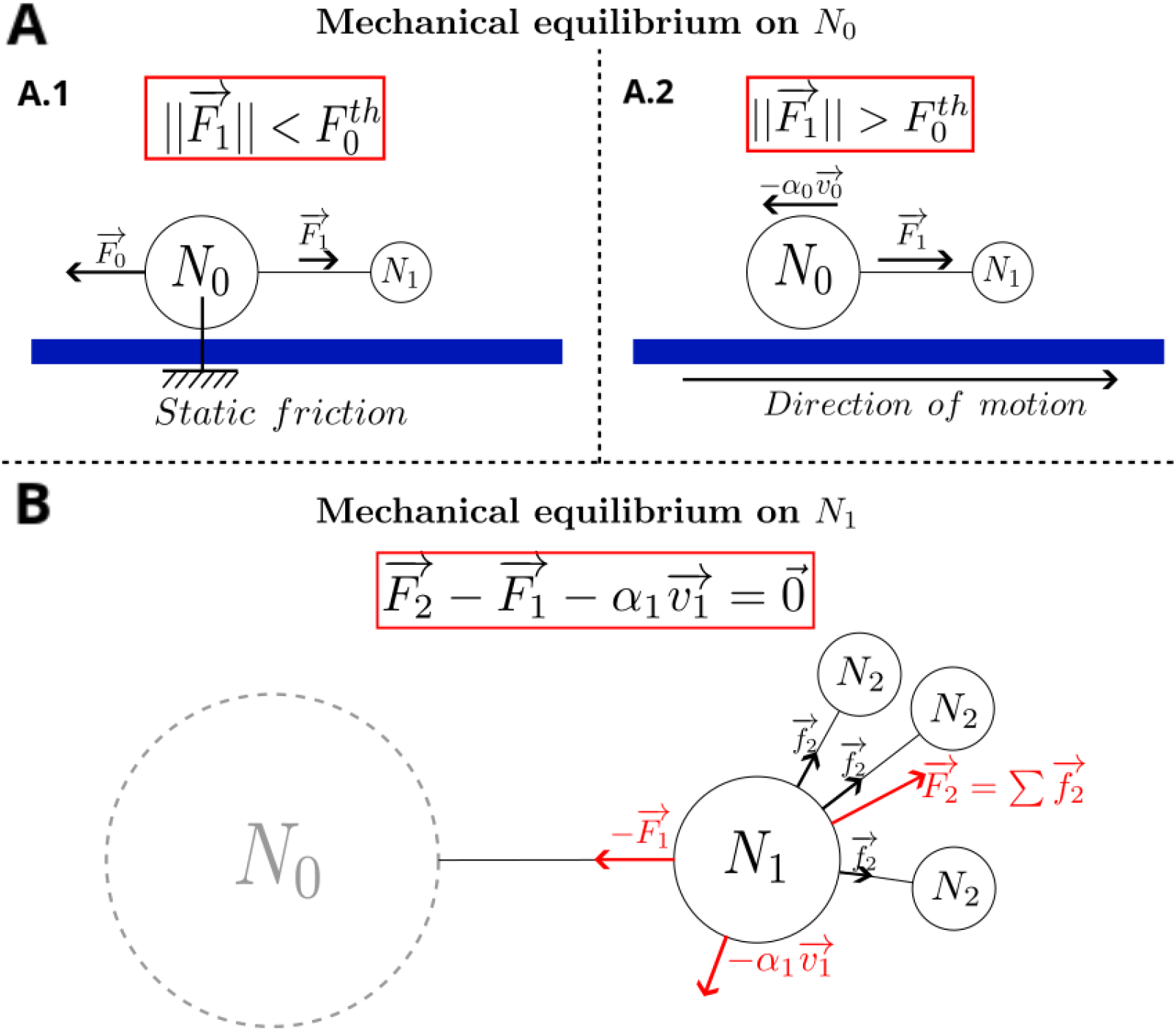
Mechanical equilibria in the model. (**A**) Force balance on the nucleus (*N*_0_) for a single adhesion. (**A.1**) The nucleus is anchored while the force norm 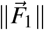 is lower than the rooting threshold 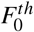. (**A.2**) The nucleus slides impaired by dynamic friction with coefficient α_0_ when 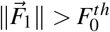. The nucleus instantaneous speed is given 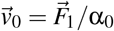 (**B**) Force balance on an adhesion site (*N*_1_). Individual *N*_2_ node exerts a force 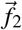 resulting in a collective force 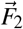. This balance applies during the nascent and protrusive phases, when *N*_1_ motion is hindered by friction (α_1_). In the contractile phase, *N*_1_ nodes are anchored and no longer generate *N*_2_ nodes.

Actin fibers are represented by the *B*_1_ branches and evolve concomitantly with the maturation of the *N*_1_ adhesions. During the first two maturation phases, *N*_1_ nodes can move relative to the substrate. Their velocity 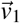 is computed under a quasi-static approximation from the balance between the force 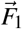 conveyed by *B*_1_ and the traction forces exerted by the *N*_2_ nodes. Interactions with the substrate are modeled through dynamic friction (Fig. 2B).

The nascent phase is exploratory, driven by the traction of multiple *N*_2_ nodes probing the environment. Due to rapid *N*_2_ turnover, the net force applied to the parent node can be approximated by a quasi-constant value *F*_2_, which both stabilizes adhesions and enables active probing. During the protrusive phase, membrane extensions characteristic of mesenchymal migration are formed. This phase is less exploratory, as the number of *N*_2_ nodes is reduced to decrease probing activity, while their traction capability is increased to further elongate and stabilize the *B*_1_ branches. In the final contractile phase, *N*_1_ nodes are fully anchored to the substrate, *N*_2_ nodes are no longer generated and *B*_1_ branches represent stress fibers that actively contract to generate traction forces sufficient to exceed the nucleus rooting threshold 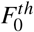.

Not all *N*_1_ nodes reach the final maturation stage, as *B*_1_ branches may rupture if the transmitted force *F*_1_ exceeds a rupture threshold. Biologically, this corresponds to insufficient integrin engagement at the adhesion site to sustain excessive traction forces. Calibration of rupture thresholds is discussed in Appendix A.

### 2.2 Maturation and mechanics of actin fibers

This section focuses on the mechanical modeling of the *B*_1_ branches and describes how a continuous force *F*_1_ progressively develops within a branch during adhesion maturation. An illustration of the buildup of force transmitted by *B*_1_ to the cell nucleus, along with the mechanical representation of each maturation phase that lasts 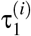 (*i* = 1, 2, 3), is shown in Fig. 3.

**Figure 3:**
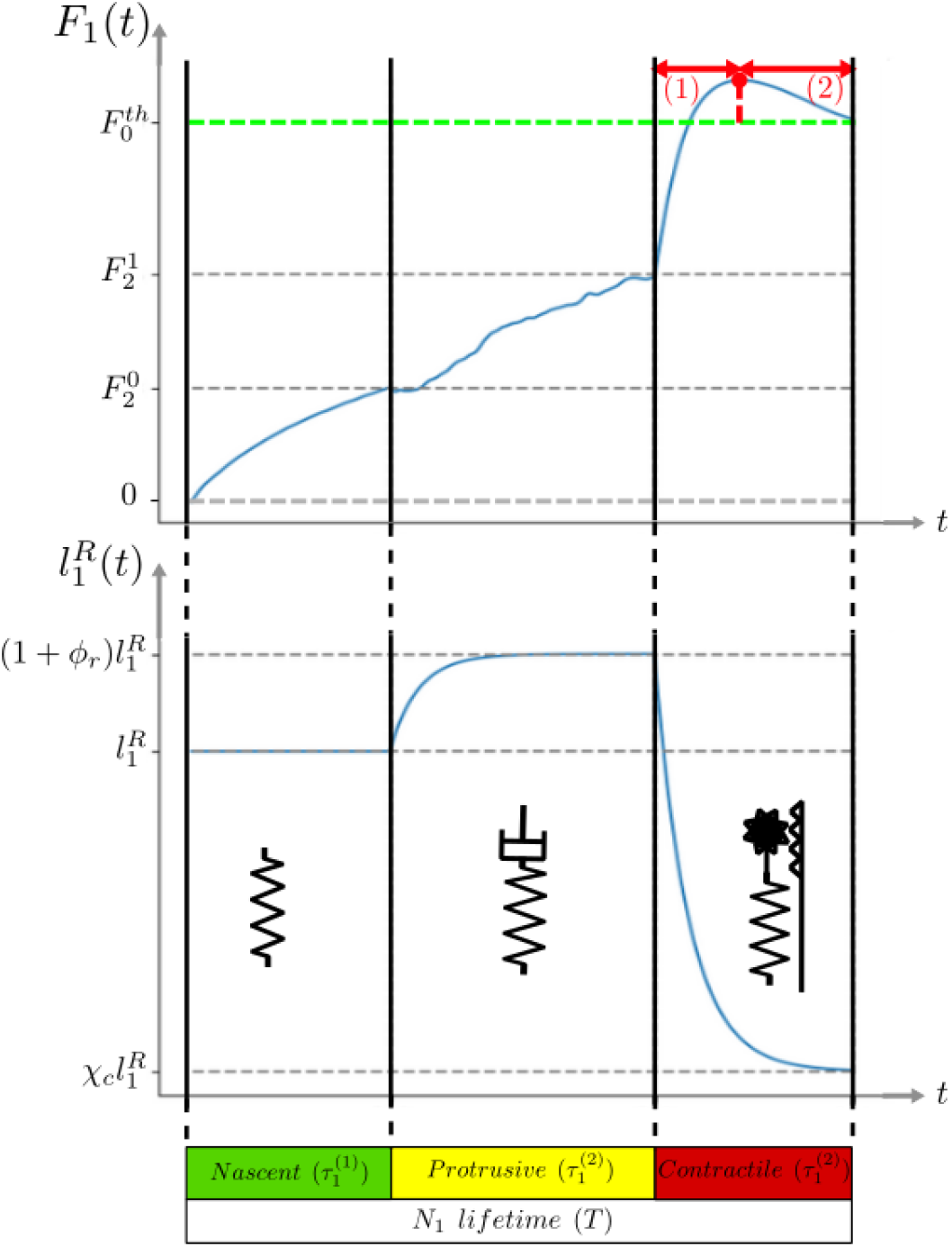
Mechanical evolution of a *B*_1_ branch during adhesion maturation. (**Top**) Representative time course of the *F*_1_ force transmitted by a *B*_1_ branch corresponding to the three maturation phases. The dashed green line indicates the nucleus rooting threshold 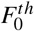 beyond which the cell moves. Two force stabilization regimes are shown: (1) the rest length decreases faster than the nucleus displacement; (2) the nucleus displacement exceeds the rest-length decrease. (**Bottom**) Time evolution of the rest length and mechanical representation of *B*_1_: spring in the nascent phase, spring–dashpot system in the protrusive phase, and actively contractile element in the contractile phase.

#### *B*_1_ Nascent phase

During the nascent phase, the *B*_1_ branch is modeled as a Hookean spring with increasing stiffness, representing the progressive bundling and alignment of actin filaments into linear structures. Stiffening is implemented through a linear increase of the apparent elastic modulus. Variability among *B*_1_ branches is introduced by drawing the rest length 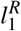 from a Gaussian distribution; this rest length is assumed constant during the nascent phase. The initial branch length is set to 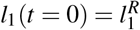, resulting in zero initial tension. Under these conditions, *N*_2_ nodes are able to elongate and reorient the branch.

#### *B*_1_ Protrusive phase

The protrusive phase is less exploratory than the nascent phase and is associated with a reduced generation frequency of *N*_2_ nodes, thereby limiting probing activity. Mechanically, this reduction decreases the collective pulling force exerted by the *N*_2_ nodes, which restricts protrusion growth and may even lead to retraction of the *B*_1_ branch. In (Chauvière et al. (2024)), protrusion growth under the sole action of the *N*_2_ nodes induces a discontinuity in the force transmitted by *B*_1_, which is not supported by experimental measurements (Du Roure et al. (2005)). To ensure a continuous force evolution, a dashpot is introduced in series with the spring. The dashpot is assumed to be fully compressed at the onset of the protrusive phase and relaxes over a characteristic time τ_*r*_ towards its unloaded steady-state length *l*_*r*_. This relaxation length is chosen as a fraction χ_*r*_ of the elongation achieved during the nascent phase:

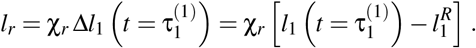

This choice reflects the assumption that a *B*_1_ branch failing to elongate during the nascent phase corresponds to unsuccessful environmental exploration and should not develop into a lamellipodium.

Dashpot relaxation is computed under the small-deformation assumption, leading to an equivalent force system represented by a single spring whose rest length evolves according to:

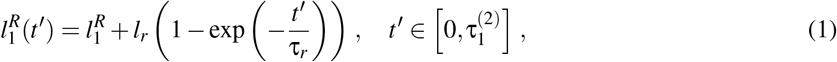

where 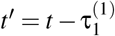 is the shifted time. The resulting evolution of rest length and force is illustrated in Fig. 3.

#### *B*_1_ Contractile phase

As in the protrusive phase, the contractile force formulation proposed in (Chauvière et al. (2024)) led to a discontinuity in the force transmitted by the *B*_1_ branch. In that framework, the force consisted of an elastic contribution and an active contractile term that grows exponentially in time, eventually dominating the elastic response and producing a length-independent force, accompanied by a force jump. Following the mechanical approach introduced for the protrusive phase, active contractility is modeled here by a time-decreasing rest length of the *B*_1_ branch. This formulation ensures force continuity and introduces a length-dependent contractile response in favor of branches that achieved significant elongation during exploration.

Specifically, the rest length is defined as:

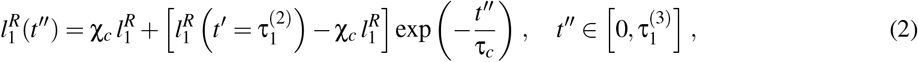

for the shifted time 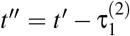. This expression ensures continuity of the rest length, and thus of *F*_1_, at the protrusive–contractile transition. It converges over a characteristic time τ_*c*_ towards a fraction χ_*c*_ of the initial branch length 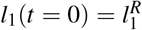, implying that larger protrusions generate stronger contractile forces. The temporal evolution of the rest length and the associated force is illustrated in Fig. 3.

### 2.3 Parametrization of the model

#### Non-dimensionalization and characteristic values

To simplify model analysis and use, all parameters are expressed in dimensionless form using biologically relevant characteristic values of time, length, force, and friction. Parameter definitions and values are summarized in Table 1.

**Table 1:** Model parameters. The column “Value” reports the parameter values used in the simulations presented in Section 3. Semicolons separate values corresponding to distinct parameters, whereas hyphens indicate lower and upper bounds of parameter ranges. Subscripts 0, 1, and 2 denote parameters associated with node levels *N*_0_, *N*_1_, and *N*_2_, respectively. Superscripts (*i*) refer to the maturation phase index.

| Symbol | Description | Value |
| --- | --- | --- |
| *Physical reference values for non-dimensionalized parameters* |  |  |
| $T$ | adhesion complex $N_1$ lifespan | 300 [s] |
| $dT$ | time step duration so that $T = n_T dT$ with $n_T = 800$ steps | 0.375 [s] |
| $L$ | mean $B_1$ branch spawn length ($l_1(t=0) = l_1^R$) | 10 [$\mu m$] |
| $F$ | nucleus $N_0$ rooting threshold ($= F_0^{th}$) | 30 [nN] |
| $\alpha$ | nucleus $N_0$ dynamic friction coefficient ($= \alpha_0$) | 0.5 [$kg.s^{-1}$] |
| *Nucleus $N_0$ - normalized parameters* |  |  |
| $F_0^{th}$ | rooting threshold | 1 |
| $\alpha_0$ | dynamic friction coefficient | 1 |
| *Adhesion complexes $N_1$ - normalized parameters* |  |  |
| $n_1$ | average number of nodes generated during $T$ | 3 – 20 |
| $\theta_1$ | exclusion angle | $\frac{\pi}{30} - \pi$ [rad] |
| $\tau_1^{(i)}$ | duration of maturation phase $i = 1, 2, 3$ (such that $\tau_1^{(1)} + \tau_1^{(2)} + \tau_1^{(3)} = 1$) | 0.3; 0.4; 0.3 |
| $\alpha_1^{(i)}$ | dynamic friction coefficients for maturation phase $i = 1, 2$ | 0.01; 0.1 |
| $F_1^{(i)th}$ | adhesion rupture threshold for maturation phase $i = 1, 2, 3$ | 0.384; 0.696; 1.5 |
| *Actin fibers $B_1$ - normalized parameters* |  |  |
| $l_1^R$ | mean initial rest length | 1 |
| $\sigma_1$ | standard deviation of initial rest length | 0.1 |
| $f_1^{(i)}$ | apparent elastic modulus at the beginning of phase $i = 1, 2$ | 0.1; 0.6 |
| $\tau_r$ | dashpot relaxation time | $0.5 \tau_1^{(2)}$ |
| $\chi_r$ | relaxation percentage of elongation of the nascent phase | 0.5 |
| $\tau_c$ | contractility characteristic time | $0.5 \tau_1^{(3)}$ |
| $\chi_c$ | percentage defining the final active branch length $l_c = \chi_c l_1^R$ | 0.01 |
| *Sensors $N_2$ - normalized parameters* |  |  |
| $n_2^{(i)}$ | average number of nodes generated during $T$ for maturation phase $i = 1, 2, 3$ | 30; 10; 0 |
| $\theta_2$ | spawn angle | $\frac{\pi}{3}$ [rad] |
| $\tau_2$ | $N_2$ lifespan | 0.01 |
| *Filipodia $B_2$ - normalized parameters* |  |  |
| $l_2$ | mean initial length | 0.4 |
| $\sigma_2$ | standard deviation of initial length | 0.01 |
| $F_2^{(i)}$ | collective pulling force in maturation phase $i = 1, 2$ | 0.3; 0.6 |

The characteristic time *T* is defined as the lifetime of an adhesion site. Experimentally measured lifetimes range from 5 to 17 min depending on cell type (goldfish fibroblasts (Kaverina et al. (2002)), PtK1 epithelial cells (Gupton and Waterman-Storer (2006))). In the simulations presented in Section 3, *T* is set to 5 min (300 s) to emphasize dynamic migration behaviours. Time is discretized into *n*_*T*_ algorithmic steps, defining a time increment *dT* such that *T* = *n*_*T*_ *dT*. The value of *n*_*T*_ is chosen to resolve the fastest processes in the model. All other time parameters are scaled by *T*.

The characteristic length *L* corresponds to the mean distance of nascent adhesions from the cell nucleus. Typical cell diameters range from 10 to 100 *µ*m (e.g., fibroblasts (Sopher et al. (2018)), epithelial cancer cells (Roux et al. (2016))). Here, *L* is set to 10 *µ*m, yielding an initial cell diameter of 20 *µ*m prior to protrusion growth. *N*_2_ nodes, representing filopodia-like structures emerging from *N*_1_ nodes, are assigned a mean length equal to 40% of *L*, consistent with their smaller size relative to lamellipodia.

The characteristic force *F* is defined as the nucleus rooting threshold 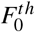, above which protrusive forces induce cell motion. Measurements using micropillar assays report values between 5 and 50 nN depending on cell type (e.g., MDCK epithelial cells (Du Roure et al. (2005)), fibroblasts and endothelial cells (Eckert et al. (2021))). A value of 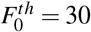 nN is used throughout this study.

The reference friction coefficient is that of the cell nucleus, denoted α_0_. It is set to 0.5 kgs^−1^ in agreement with estimated value measured by surface unit in (Chelly et al. (2022)). This yields a minimum instantaneous nuclear velocity 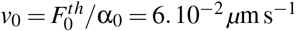 (216 *µ*m h^−1^; see Eq. (A.6)). As shown in Section 3, *in silico* cells are motile for 10–70% of the simulation time, resulting in average speeds between 20 and 150 *µ*m h^−1^. While the upper bound is relatively high, this range includes experimentally reported values, from ∼ 15 *µ*m h^−1^ for CHO cells (Harms et al. (2005)) to ∼ 60 *µ*m h^−1^ for fibroblasts (Pham et al. (2016)).

#### Key mechanical conditions for cell movement

Rather than performing an exhaustive numerical exploration of the full parameter space, our analysis focuses on identifying conditions that reproduce experimentally observed features of cell migration. This section summarizes the key mechanisms governing the onset of cell motion in the *in silico* model. The corresponding analysis is provided in Appendix A.

Cell motion occurs when the net force acting on the nucleus *N*_0_ (i.e., the sum of forces transmitted by all *B*_1_ branches) exceeds the rooting threshold 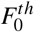 separating static and dynamic friction regimes (Fig. 2A). For analytical clarity, a single adhesion site *N*_1_ is considered. In this case, the force *F*_1_ transmitted by the corresponding *B*_1_ branch can be estimated from force balance at *N*_1_ (Fig. 2B). Because the mechanical properties of *B*_1_ branches evolve during maturation, the temporal evolution of a single force *F*_1_ is analyzed across the three maturation phases. This analysis yields parameter conditions required to reproduce biologically consistent behaviour, defined by the following criteria:

- a single adhesion site must succeed in maturing without breaking – conditions (A.1) and (A.2) in Appendix A;
- protrusion growth must occur during the first two phases of the maturation process – condition (A.4);
- cell translocation must occur only during active actin contraction in the final maturation phase – conditions (A.1), (A.2) and (A.5).

These conditions provide a mechanical calibration of the model. The parameter values used in numerical simulations are listed in Table 1. The time evolution of a single protrusive force *F*_1_ acting on the nucleus and driving active cell translocation and cell movement is illustrated in Fig. 3.

## 3 Results

### 3.1 Cell morphology

A simplified scenario in which a single adhesion is sufficient to trigger cell motion was analyzed and used to calibrate the model. Under this calibrated parameter set, multiple adhesions are subsequently allowed in order to explore the range of cell morphologies that the model can reproduce and to assess their influence on cell dynamics.

Cell morphology is defined here as the characteristic shape maintained by a cell over time. It is therefore characterized by the average number of *N*_1_ nodes and by the degree of synchronization of their maturation stages. These properties are controlled by two key parameters:

- θ_1_: an exclusion angle preventing the emergence of a new *N*_1_ node in the vicinity of an existing one. This parameter controls the range of shapes that a cell can display (see Fig. 1);
- *n*_1_: the average number of *N*_1_ nodes generated during the characteristic time *T*, defining the *N*_1_ generation frequency *n*_1_/*T*. This parameter also governs the synchronization of *N*_1_ maturation phases. Indeed, large values of *n*_1_ favor the rapid formation of multiple adhesions (when geometrically permitted by θ_1_) leading to concurrent maturation of the nodes.

Over the characteristic time *T*, the average number of *N*_1_ nodes around the cell is well approximated by 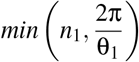 as illustrated in Fig. 4. When θ_1_ is small, many potential sites are available for *N*_1_ emergence, and the generation frequency primarily controls the number of adhesions. Conversely, for large θ_1_, the number of available sites is limited, and spatial constraints dominate. This quantity thus provides a concise descriptor of *in silico* cell morphology. Three distinct regimes emerge, leading to different cell morphologies as illustrated in Fig. 4, and associated dynamics (described in the next section) :

**Figure 4:**
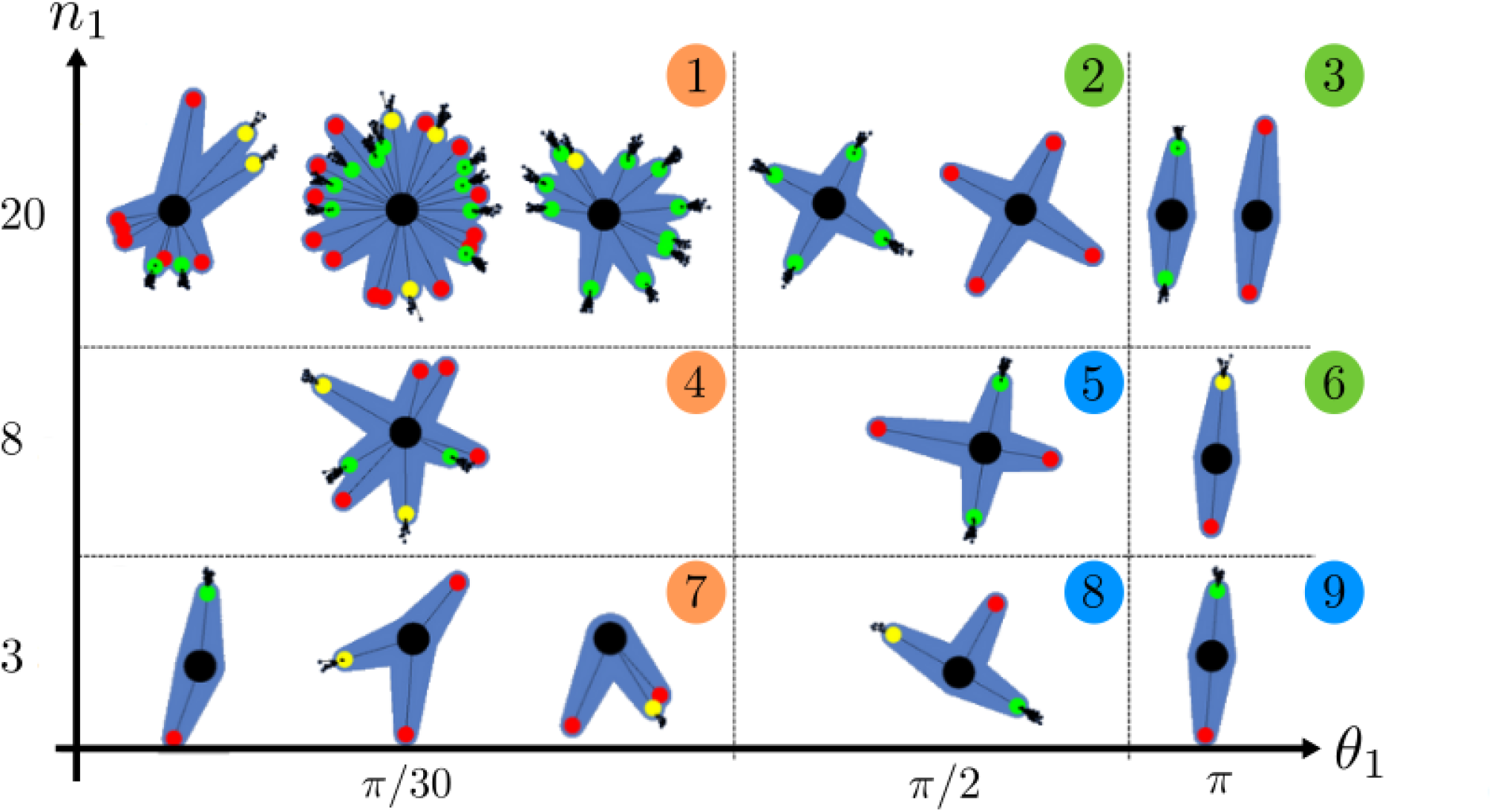
Illustration of *in silico* cell morphologies for different values of the exclusion angle θ_1_ and *n*_1_ defining the *N*_1_ generation frequency *n*_1_/*T*. Colors refer to the three regimes, whereas numbers correspond to different examples, as described in text.

1. 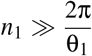. In this regime, the exclusion angle dominates and the cell exhibits an almost time-independent shape, corresponding to a stable morphology. Most admissible *N*_1_ nodes are generated within a short time window and thus mature nearly synchronously. As a result, the contraction forces generated by opposing adhesions tend to balance each other, strongly impairing cell motion. Numerical examples (2, 3, and 6) in Fig. 4 illustrate this regime.
2. 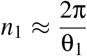. Here, the cell reaches its maximal number of adhesions when the earliest formed node ends its contractile phase. Adhesion maturation is less synchronized, giving rise to a cyclic contraction pattern. When an adhesion disappears due to aging, a single site becomes available for the emergence of a new node. This regime corresponds to intermediate stability of cell morphology and moderate cell motility. Numerical examples (5, 8 and 9) in Fig. 4 illustrate this behaviour.
3. 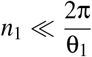. In this case, the *N*_1_ generation frequency determines the average number of adhesions, and cell morphology is unstable over time. Reduced competition between contracting adhesions results in enhanced cell motility. Numerical examples (1, 4, and 7) in Fig. 4 correspond to this regime.

### 3.2 Cell dynamics and trajectory analysis

Cell dynamics and the resulting migratory features are analyzed depending on cell morphology, *i*.*e*., as function of the parameters θ_1_ and *n*_1_. Variations in these two parameters are indeed the primary source of qualitative changes in cell dynamics. In contrast, the remaining parameters (mechanical properties, size, and characteristic times) mainly act as scaling factors, affecting quantities such as traveled distance, migration speed, force magnitude, or characteristic metrics including the diffusion coefficient. The analysis is performed accordingly using fixed values for all other parameters (see Table 1) chosen to ensure effective cell motion. In particular:

- a single *N*_1_ node is able to generate a traction force exceeding the nucleus rooting threshold;
- adhesion rupture forces are calibrated based on the analysis summarized in Section 2.3 and detailed further in Appendix A.

In this section, the range of cell dynamics produced by the model is characterized, with particular emphasis on the three morphological regimes identified in the previous section. This analysis establishes a link between cell morphology and migratory dynamics, which may be interpreted as representative of distinct cell types, and validates the parameter calibration introduced in Section 2.3 as a mean to reproduce diverse and biologically relevant cell morphologies and migration patterns.

#### Cells migrate as a persistent random walk

Two limiting regimes are classically considered when analyzing trajectories: ballistic motion, in which objects move along straight paths, and diffusive motion, characterized by random displacement. As emphasized in (Pawluchin and Galic (2022)), cell migration typically lies between these two extremes and is well described as a persistent (biased) random walk, which can be modeled using run-and-tumble or Lévy flight frameworks. This migratory behaviour is characterized by a transition from ballistic motion at short time lags to diffusive motion at longer timescales. Such a transition can be identified by computing the mean-squared displacement (MSD) ⟨*x*^2^⟩(δ*t*) ∝ δ*t*^α^. On a logarithmic scale, the MSD exhibits therefore a linear increase with slope α = 2 in the ballistic regime and α = 1 in the diffusive one. This change of slope is characterized by the so-called directional persistence time.

As a first observation, trajectories obtained from the simulations (Fig. 5) qualitatively resemble experimentally observed *in vitro* trajectories (Brückner and Broedersz (2024); Peschetola (2011)) as well as previously reported numerical simulations (Scianna et al. (2013)). To start a more quantitative study, the analysis of the MSD is performed using bootstrapping applied to individual cell trajectories and subsequently averaged over cells sharing the same parameter set. Note that, although bootstrapping may be questionable due to the non-Markovian nature of the simulations — stemming from the maturation dynamics of adhesion nodes — the total simulation time is approximately 50 times larger than the characteristic timescale of the system. As a result, memory effects are negligible and no significant differences were observed between analyses performed with or without bootstrapping, aside from improved smoothness of the curves when bootstrapping is employed. The behaviour of the MSD over time is shown on Fig. 6A : once cells are able to generate sufficient traction to move, they systematically display a persistent random walk; this allows to detect biologically inconsistent behaviours - referred to as “pathological” simulations - by the shape of their MSD curve as illustrated in the insert of the figure. More precisely, the short-time ballistic behaviour arises from the contraction of a single *B*_1_ branch, which drags the cell nucleus in a preferential direction. After multiple contraction events, the trajectories progressively lose directional bias and exhibit diffusive behaviour. The transition between the two regimes is then characterized by the duration of the contractile phase, which provides therefore a first approximate of the directional persistence time.

**Figure 5:**
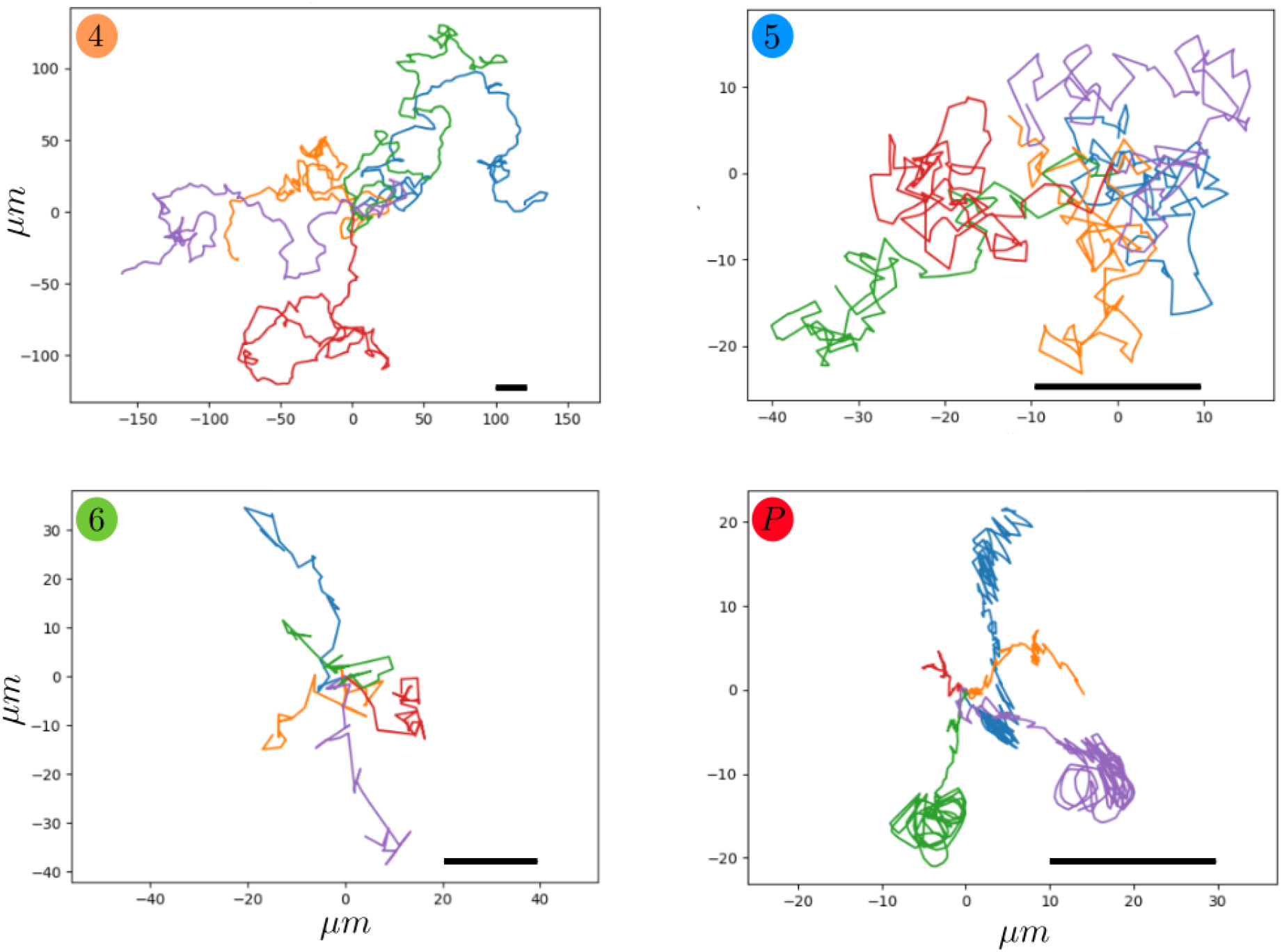
*In silico* cell trajectories of five cells over 40000 time steps (≈ 4 *h*). Panels 4,5 and 6 refer to the morphology chart indexation (Fig. 4). Box (P) is an example of a “pathological” case for 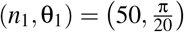 and high adhesion rupture thresholds yielding unbreakable adhesions.

**Figure 6:**
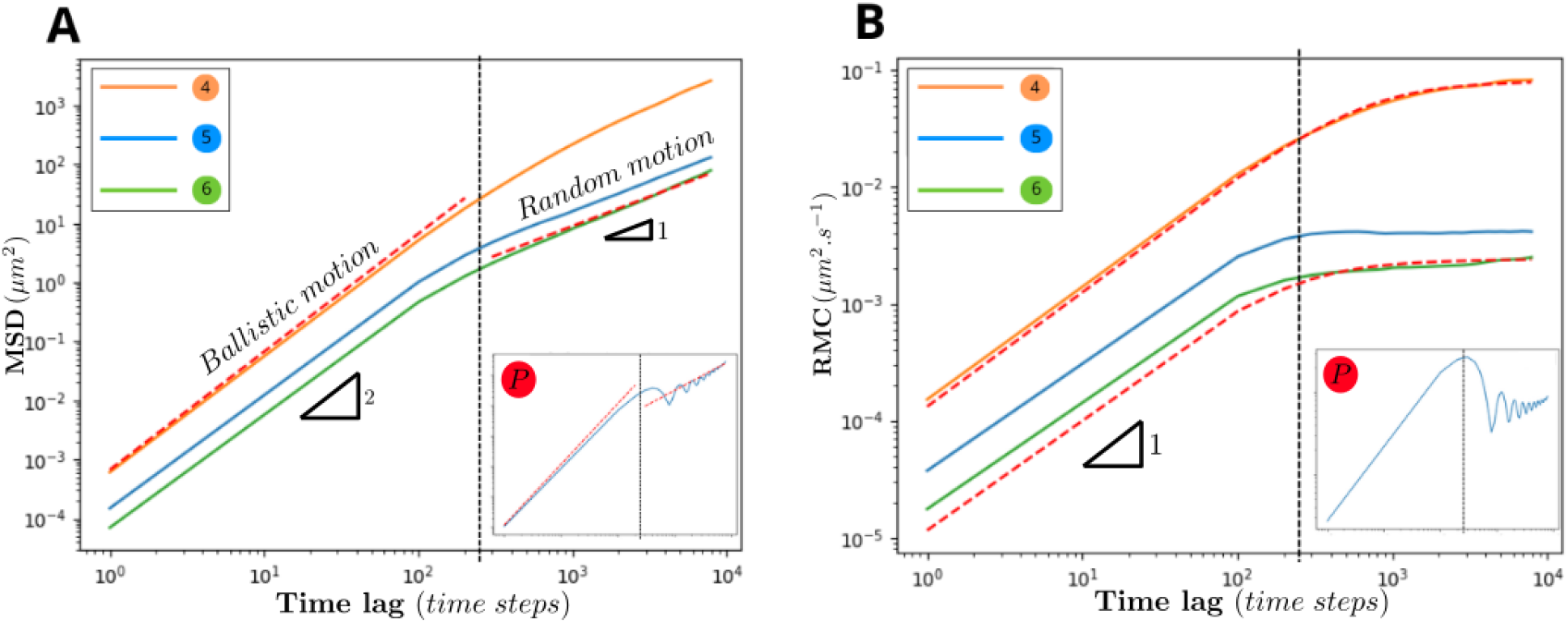
Trajectory analysis of cases 4, 5 and 6 on Fig. 5 - Insert corresponds to the pathological case. **A** : Mean squared displacement (log-log axis). The red dashed lines are affine functions with slope 2 for ballistic motion and 1 for random motion. **B** : Random motility coefficient. The red dashed lines are fitted random motility coefficients based on theoretical persistent random walk. On both plots, the dark dashed line indicates the duration of the contractile phase 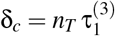 in time steps.

A precise characterization of the movement is obtained by considering the theoretical modeling of a persistent random walk in two dimensions (see (Brückner and Broedersz (2024); Dickinson and Tranquillo (1993)) for details). The MSD is then given by ⟨*x*^2^⟩(δ*t*) = 2*S*^2^*P*(δ*t* − *P*(1 − *e*^−δ*t*/*P*^)) where *S* is the root mean squared speed and *P* is the directional persistence time. Accordingly, the random motility coefficient (RMC) - defined by ⟨*x*^2^⟩(δ*t*)/4δ*t* - increases linearly at short time lags with a slope *S*^2^/4, and reaches a plateau equal to *S*^2^*P*/2 at long time lags. This behaviour is recovered in the simulations as shown on Fig 6B, and allows for the determination of *S* and *P* in each cases. In particular, for cells with an exclusion angle θ_1_ = π - case 6 - the fitted values are *S* = 1.810^−2^ *µm*.*s*^−1^ and *P* = 39.4 *s*, which is shorter than the duration of the contractile phase (90 *s*). This reduced persistence arises from counteracting forces generated by adhesions on opposite sides of the cell. In contrast, for smaller exclusion angles - case 4 - larger values are obtained (*S* = 5.610^−2^ *µm*.*s*^−1^ and *P* = 150 *s*), indicating faster cells with enhanced directional persistence extending beyond a single contraction event, as discussed in the following section. Theoretical random motility coefficient curves corresponding to these values of *S* and *P* are shown as red dashed lines in Fig. 6B.

#### Cell directionality

Although the mean-squared displacement provides a useful classification of migratory regimes, it is not sufficient to discriminate between distinct cell phenotypes. Indeed, different underlying mechanisms may produce similar MSD profiles, as illustrated by the variety of trajectories in Fig. 5 yielding comparable MSD curves that differ primarily in their random motility coefficient.

As a first indicator of directionality, tortuosity was computed as the ratio of the total distance *d*(*t*) traveled up to time *t*, to the distance 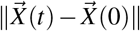 between the cell position at time 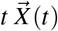 and its initial position 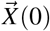. As shown in Fig. 7A, the tortuosity exhibits a transient stabilization slightly below 0.2, in agreement with values reported in Stéphanou et al. (2008), for fibroblast migration. This stabilization is however transitory as, over long timescales, cell trajectories are driven by randomness and the tortuosity ultimately converges towards zero.

**Figure 7:**
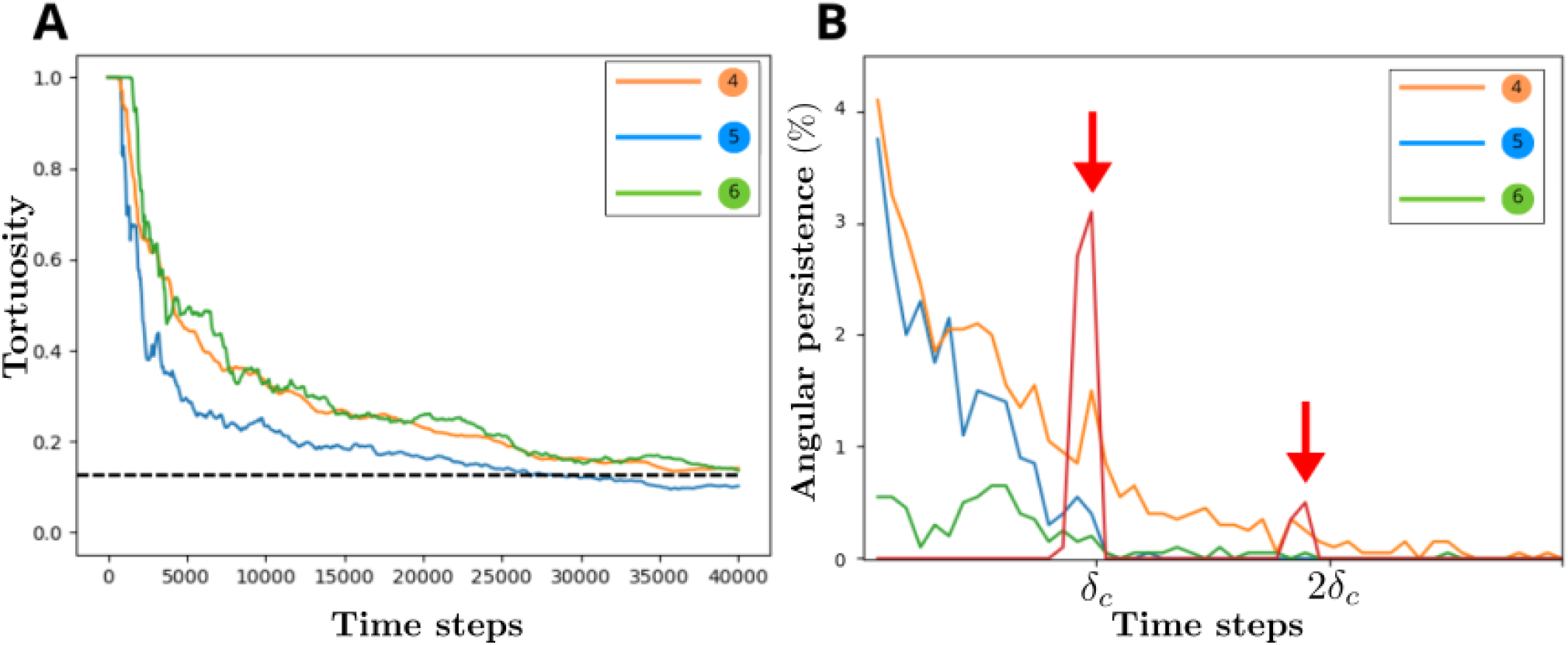
Directionality indicators of cases 4, 5 and 6 on Fig. 5. **A** : Average tortuosity of cell trajectories. The dark dashed line marks the stabilization value. **B** : Angular persistence during cell movement. The control case of a cell growing a single branch (θ_1_ = 2π) is shown in red to highlight the peaks (red arrows) occurring at once and twice the contractile phase duration δ_*c*_.

Cell directionality was further characterized by the angular persistence, *i*.*e*. the percentage of time consecutively spent by a cell in a direction of movement that remains in a cone of fixed angle (that was arbitrarily fixed to the value π/3 for illustrative purpose). This second indicator is obtained thanks to the unit-velocity autocorrelation function 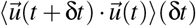 with 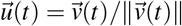 the unit vector providing the direction of the cell movement at time *t*. Specifically, this indicator corresponds to the number of consecutive time steps during which the cell velocity direction remains within the cone of fixed angle, divided by the total duration of the cell trajectory. Time steps corresponding to stationary phases were excluded; When a cell resumes motion after a pause, the persistence is conserved if the new velocity direction lies within the cone defined by the last nonzero velocity vector; otherwise, it is reset to zero. Results are shown in Fig. 7B. First, as time steps corresponding to immobile phases are excluded from this evaluation, not surprisingly the angular persistence displays low values because stationary phases account for approximately 50% to 90% of the total simulation time (as evidenced in Fig. 8B). Second, a control case for cells restricted to forming a single branch at a time (*i*.*e*., for θ_1_ = 2π) is presented : peaks of angular persistence are observed at times corresponding to once and twice the contractile phase duration, respectively associated to one and two consecutive contractions of the only branch of the cell. Consistent with the chosen uniform angular distribution of *N*_1_ node emergence (leading to two successive contraction events occurring within a cone of angle π/3 with probability 1/6), the second peak is six times smaller than the first one at δ_*c*_.

**Figure 8:**
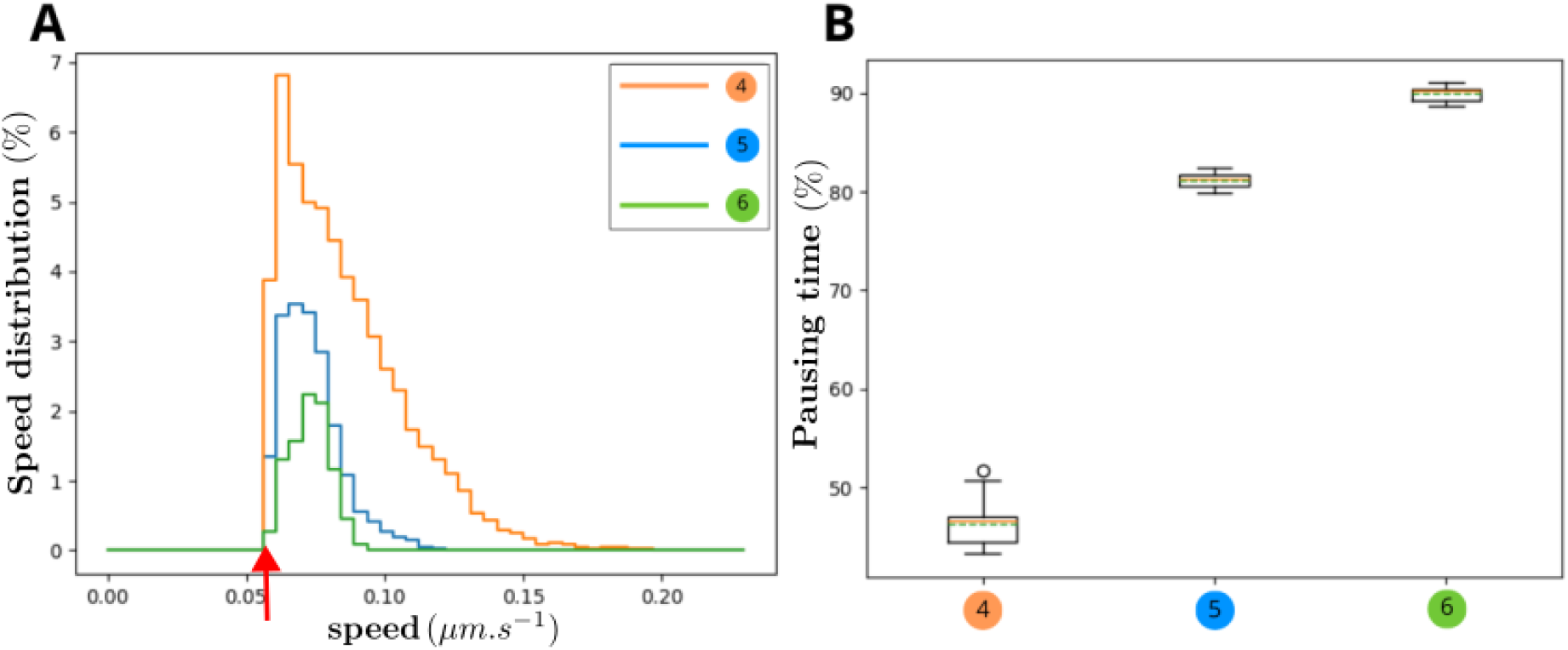
Cell speed and pausing time for cases 4, 5 and 6 on Fig. 5. **A**: Distribution of cell speed (null speed excluded). The red arrow indicates the theoretical minimal instantaneous cell speed, 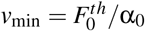. **B**: Pausing time expressed as the ratio of immobility to total trajectory time.

For the parameter sets used in this study, cell movement is performed with low persistence – *i*.*e*., observed persistence times below δ_*c*_ – causing trajectories to curl and underlying the diffusive behaviour observed at long timescales. Yet a jump around δ_*c*_ can be observed, consistent with the slope rupture of MSD curves. Nevertheless, for small exclusion angles – case 4 – cells are able to maintain directional persistence over longer periods, in some cases exceeding three times the contraction phase duration. This behaviour arises from successive contraction events occurring on the same side of the cell, leading to repeated nucleus translocation in a preferred direction. As a consequence, branches on the opposite side experience increased tension, eventually leading to rupture, while adhesions on the front side persist and mature, reinforcing directional motion.

The model is intended as a baseline framework for subsequent investigations of cell interactions with deformable substrates, where migration may be biased by mechanical cues sensed through the probing capacity of *N*_2_ nodes. On a rigid substrate, the emergence of a persistent random walk with limited directional memory is expected, as no external asymmetry is available to guide migration. In particular, no explicit polarization maintenance mechanism is implemented, as the long term objective is to investigate the emergence of directed motion solely from mechanical interactions. In this context, the present trajectory analysis serves as a validation step for the migration model and provides a reference state for comparison with future results obtained on deformable substrates.

#### Cell speed and migration time

To conclude the analysis of the model, cell speed distributions were compared across simulations and shown in Fig. 8A. A clear trend emerges: decreasing the exclusion angle θ_1_ leads to higher cell speeds. The increase in speed observed for small exclusion angles can be explained by two complementary mechanisms. First, several *N*_1_ nodes may contract in similar directions, leading to an accumulation of traction forces on the nucleus that largely exceeds the rooting threshold. This situation is typically observed for low *N*_1_ emergence frequency combined with small exclusion angles, which occasionally allow multiple adhesions to form in close proximity without an opposing force on the opposite side of the cell. Second, for high *N*_1_ emergence frequency, the cell may remain stationary due to opposing contraction forces acting on the nucleus. When one adhesion disappears the force balance is suddenly broken, releasing the load and propelling the cell at a speed higher than the minimal value : as the force in the remaining *B*_1_ branches continues to build up and exceeds the rooting threshold at the time the opposing load is released.

The minimal instantaneous cell speed is set by the model parameters and is given by 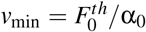. This property facilitates calibration of the model to experimental data corresponding to specific cell types: the nucleus rooting threshold 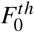 and the effective friction coefficient α_0_ can be adjusted to reproduce a given speed. This speed is however an instantaneous quantity, whereas experimentally reported speeds are typically averaged over finite acquisition intervals. To recover average migration speeds, it is therefore necessary to account for the pausing behaviour of cells illustrated in Fig. 8B.

For large exclusion angles, cells remain stationary for up to 90% of the time. This fraction decreases as the exclusion angle is reduced, reaching slightly less than 50% for θ_1_ = π/30. An exception to this trend occurs for exclusion angles larger than π (not shown) for which cells can only form a single adhesion at a time: In this case, cell motion is restricted to the contraction phase, which represents 30% of the total time for the chosen parameter set. Large exclusion angles restrict the number of simultaneous *N*_1_ nodes, promoting configurations in which contraction forces act in opposing directions. This mutual force cancellation decreases the duration over which the net force on the nucleus exceeds the rooting threshold. Consequently, the primary determinant of the decrease in pausing time with decreasing exclusion angle is the number of available directions in which the cell can generate traction forces. The larger the number of available directions, the lower the probability that all traction forces are mutually canceled. This effect is apparent in the cell trajectories shown in Fig. 5, where highly motile cells – case 4 – display more curved paths, whereas cells exhibiting prolonged pauses – case 6 – tend to move along straighter trajectories.

## 4 Discussion

In this work, a mechanistic model of mesenchymal cell migration is introduced. It is designed to capture the emergence of motility, persistence, and polarization from intracellular traction forces alone. By explicitly modelling actin-driven protrusions (*B*_1_ branches) as force-generating elements anchored to the substrate, the framework links cell-scale migration patterns to well-defined mechanical rules acting at the level of adhesions and the nucleus. A central result of this study is that, in the absence of external cues, the model robustly reproduces a persistent random walk, a hallmark of mesenchymal migration observed across experimental and numerical studies. This behaviour emerges without prescribing polarization or directional memory, thereby validating the model as a mechanically grounded reference case for unbiased cell migration.

Beyond reproducing qualitative migratory patterns, the model demonstrates quantitative flexibility. By tuning a limited set of parameters, it can be calibrated to match experimentally relevant cell properties such as migration speed, traction force levels, and pausing behaviour. The mechanical analysis of force balance within branches further allowed us to delineate regions of the parameter space that support migration from those leading to immobile states – either due to insufficient force generation or excessive adhesion stability. This systematic characterization strengthens the interpretability of the model and clarifies the mechanical conditions under which mesenchymal-like motility can arise.

While these results support the relevance of the proposed framework, several limitations should be acknowledged. First, the model focuses on a single cell migrating on a static, rigid substrate. As a consequence, key feedback mechanisms known to influence migration – such as substrate deformation, stiffening, and force redistribution – are intentionally neglected at this stage. Second, the representation of adhesions is coarse-grained: individual *N*_1_ nodes should not be interpreted as biological focal adhesions, but rather as subunits of adhesion complexes whose collective behaviour mimics effective adhesion sites. This modelling choice explains why increased numbers of *N*_1_ nodes promote persistence and motility in the present framework, in apparent contrast with experimental observations reporting an optimal focal adhesion number in 3D environments (Fraley et al. (2010)). In addition, the model is restricted to 2D migration, which differs fundamentally from 3D motility where confinement, steric hindrance, and matrix porosity play a dominant role (Yamada and Sixt (2019); Caswell and Zech (2018)). These simplifications are deliberate and reflect the objective of isolating mechanical principles while keeping the computational cost compatible with future extensions.

Other modelling assumptions may require reassessment in more complex environments. For instance, the criterion allowing a *B*_1_ branch to mature into a contractile state based on its elongation was deactivated here (with respect to the original model), as mechanical analysis showed it to be either always satisfied or never reached for the parameter ranges explored. On a deformable substrate, however, traction-induced displacement of adhesion points would effectively shorten branches, reducing transmitted forces and potentially preventing maturation. In that context, branch-length–dependent maturation could become a critical mechanism for selectively reinforcing adhesions on stiff regions, without introducing additional ad hoc rules. Similarly, while introducing biased adhesion nucleation could artificially increase persistence, this option was deliberately avoided to ensure that polarization arises from mechanical interactions rather than imposed spatial correlations.

Taken together, the present model provides a minimal and mechanically grounded reference framework for cell migration in the absence of external cues. Its ability to reproduce persistent random walk statistics solely from intracellular force generation and adhesion dynamics makes it particularly suitable as a control case. The natural next step is to couple this migration model with a deformable extracellular matrix, enabling cell-generated forces to locally modify substrate architecture and stiffness. In such a context, mechanical feedback alone is expected to bias migration, giving rise to durotactic responses without imposing additional polarity rules. Ultimately, this approach will allow investigation of how cell-induced ECM deformations, including fiber reorientation and stiffening, influence migratory trajectories and facilitate encounter processes between migrating cells, a key aspect in the context of anastomosis.

## 5 Acknowledgments

We thank Daria Tsvirkun from LIPhy laboratory for providing the cell image of Fig. 1. This work is part of the MoDyMecA project funded by ANR (ANR-23-CE45-0026 MoDyMecA). This manuscript benefited from the use of OpenAI’s ChatGPT (GPT-5) to improve English clarity and correctness.

## 6 Supplementary section

The code used to generate the simulations along with videos of *in silico* cells in different parameter sets and additional comparative plots are provided in the supplementary section available at : https://gricad-gitlab.univ-grenoble-alpes.fr/timc-bcm/cellmotility_articleversion.

## Statements and Declarations

### Conflict of interest

The authors declare no conflict of interest.

## A Model Analysis and calibration

In this section the aim is to identify regions of the parameter space where cell movement occurs in a biologically consistent manner. Cell movement occurs in the model when the net force on the nucleus (*i*.*e*. the sum of the forces in *B*_1_ branches acting on the central node *N*_0_) exceeds a rooting threshold. Thus the understanding of cell movement requires to know the mechanical behaviour in those *B*_1_ branches. Therefore the force balance on a single adhesion complex *N*_1_ is analyzed and the mechanical dynamics of the single associated *B*_1_ branch over time (during the three maturation phases) is evaluated.

When an adhesion complex *N*_1_ is allowed to move (*i*.*e*. when non-mature, which corresponds to the first two maturation stages), it velocity 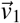 results from the following balance of forces on *N*_1_ :

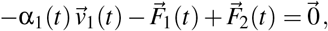

where α_1_ is the time-dependant friction coefficient of 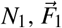 the elastic restoring force in *B*_1_ and 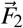 the collective pulling force of *N*_2_ nodes (see Fig. 2B). The movement of *N*_1_ serves to lead the force balance at equilibrium, which is eventually given by the scalar equality resulting from the projection on the *N*_0_*N*_1_-axis :

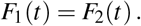

This will be the base ingredient of our calibration process. Note that in the following sections, the maturation stage notation ^(*i*)^ is often omitted to reduce the amount of notation when no confusion is possible.

### 1. Nascent phase (1)

Main hypotheses in the nascent phase :

1. The *B*_1_ branch is a spring of constant rest length 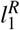 leading to a restoring force 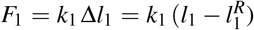, which we rewrite as *F*_1_ = *f*_1_ ε_1_ with corresponding strain 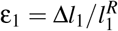 Here *f*_1_ has the dimension of a force but can be seen as an “apparent elastic modulus” given by the product of the Young modulus by a given branch section area.
2. The stiffness coefficient *k*_1_ linearly increases over time during this phase, which results in the linear increase of *f*_1_ from 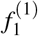 to 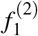 to account for fibers reinforcement.
3. Similarly, the friction coefficient α_1_ linearly increases over time from 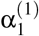 to 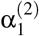 to account for cell adhesion molecules recruitment at the adhesion site.
4. The *B*_1_ branch is not under tension when created, *i*.*e*. the restoring force *F*_1_(*t*) = *f*_1_(*t*) ε_1_(*t*) is set to zero at *t* = 0 by imposing 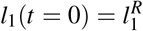. (See Fig. 3). The tension in the branch eventually occurs due to the presence of *N*_2_ nodes.
5. The *n*_2_ extremities *N*_2_ of *B*_2_ branches are anchored to the substrate to generate a net pulling force 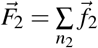 on *N*_1_ with intensity *F*_2_ in the *B*_1_ branch direction.
6. Each *B*_2_ branch is a spring of constant stiffness *k*_2_ initially under maximal tension by verifying Hooke’s law 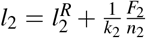. As *N*_2_ are anchored whereas *N*_1_ is allowed to move, the force intensity *f*_2_(*t*) in each *B*_2_ decreases over time.

#### 1.1 Calibration of *F*_2_ with respect to 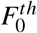

When a single branch *B*_1_ acts on the nucleus *N*_0_, the nature of the nascent phase of the associated *N*_1_ requires that *F*_1_ does not reach the critical value 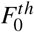. Indeed, in its initial phase, the adhesion complex is not mature enough to lead to nucleus translocation. Therefore, for biological consistency of this initial phase, the intensity *F*_1_ of the force in the *B*_1_ branch has to always be lower than the critical value 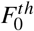, *i*.*e*. 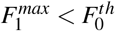. Three scenarii of interest can be observed in our simulations:

a. the collective pulling force *F*_2_ generates the elongation of *B*_1_ (hence the increase of *F*_1_) not fast enough to reach equilibrium;
b. the equilibrium is reached during the nascent phase;
c. the variation over time of α_1_ and *f*_1_ leads to a non-monotonic behaviour of *F*_1_ with maximal theoretical value 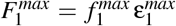, where 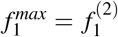 and the maximal strain is achieved when all *N*_2_ branches are under maximal tension (*i*.*e*. at their appearance time), which leads to 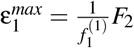. Therefore the estimated value 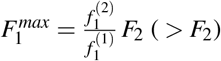 is obtained.

To select *F*_2_ so that 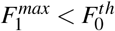, the following condition is chosen :

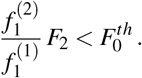

#### 1.2 Calibration of 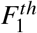 with respect to *F*_2_

Adhesion can experience rupture if the conveyed force *F*_1_ exceeds the rupture threshold 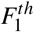, leading to the death of the node. But *N*_1_ nodes should be able to mature in the nascent phase, and thus not to systematically break adhesion to the substrate. Therefore the condition 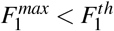 is required to provide admissible values of 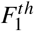, which leads to the condition

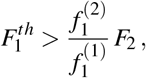

thanks to the estimated value of 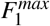. During this phase, adhesion sites have to break more easily than nucleus translocation, which leads to another condition

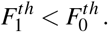

#### 1.3 Conditions on mechanical parameters

When regrouping the above conditions, mechanical parameters (written with the indication of the corresponding maturation phase for the reader clarity) have to be chosen such that

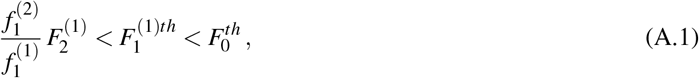

to insure relevance of the biological behaviour of the *in silico* cell.

### 2. Protrusive phase (2)

Main hypotheses in the protrusive phase :

1. The *B*_1_ branch is a spring of constant stiffness given by the apparent elastic modulus 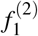 (reached at the end of the previous phase), in series with a dashpot. (See Fig. 3).
2. The dashpot is initially compressed and relaxes to its characteristic length *l*_*r*_ in time τ_*r*_.
3. The spring-dashpot system is equivalent to a single spring with time-dependent rest length - see Equation (1) in main text - starting from 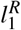 and converging over time towards 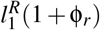 where 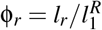.
4. The friction coefficient is assumed constant at the value 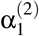 (reached at the end of the previous phase).
5. All *n*_2_ nodes *N*_2_ and associated *B*_2_ branches maintain the same mechanical behaviour as described in the previous phase.

In this phase the pulling force exerted by the *N*_2_ nodes exhibits a transient regime to evolve from 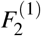 to 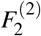. This transitory regime lasts a maximal time corresponding to the life span τ_2_ of *N*_2_ nodes, which is short relatively to the maturation phase duration 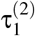. In the meantime, the dashpot starts relaxing, reducing exponentially the magnitude of *F*_1_. As in the nascent phase, the adhesion complex is still not mature enough and should break more easily than nucleus translocation, which leads again to the condition 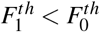. Then two scenarii of interest can be observed in our simulations:

a. If 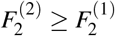 then the *N*_2_ nodes are mechanically more active than in the nascent phase and drive the protrusion growth (*F*_2_ > *F*_1_ during the whole protrusive phase) of the *B*_1_ branch (whose dashpot is therefore not required but plays an important role in the protrusion mature size). In this case, in order to allow *N*_1_ nodes to remain adherent (*i*.*e*., no rupture), the condition 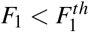 has to be insured for all times, which is always satisfied when imposing 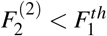.
b. If 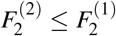 then the *N*_2_ nodes are mechanically less active than in the nascent phase and the protrusion growth shall be carried by the internal mechanics of the *B*_1_ branch (*i*.*e*. the dashpot yielding the time-varying rest length is required in this case - see below). Then *F*_1_ shall balance the new value of *F*_2_, thus decrease from 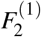 to 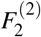. Adhesion break should therefore be avoided by imposing 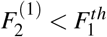.

#### 2.1 Conditions on mechanical parameters

When regrouping the conditions above, mechanical parameters have to be chosen such that

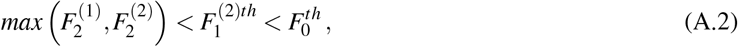

to insure relevance of the biological behaviour of the *in silico* cell.

Remark that protrusion growth in case (b) requires conditions on the dashpot properties and cannot be achieved arbitrarily. To support this remark, let us assume that the friction force is negligible to reach the mechanical equilibrium of an *N*_1_ node (in each of its nascent and protrusive phase), given by the balance

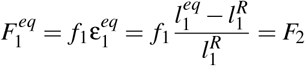

and thus by the equilibrium length

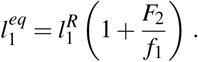

Protrusion growth during the protrusive phase requires therefore that 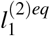-the equilibrium length at the end of phase (2) - is larger than 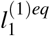-the equilibrium length at the end of phase (1) - to insure that the length of the *B*_1_ branch increases, where

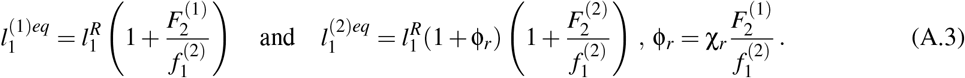

Therefore 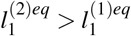 is equivalent to choosing a minimal value of χ_*r*_ such that:

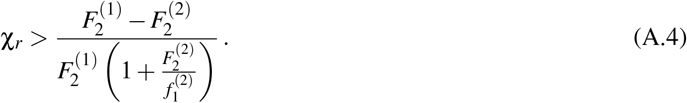

Whereas this condition is always satisfied in case (a), one must expect this condition to be relevant in order to observe protrusion growth in case (b).

### 3. Contractile phase (3)

Main hypotheses in the contractile phase :

1. The *B*_1_ branch is a spring of constant stiffness associated with the apparent elastic modulus 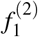 (in continuity with the previous phase).
2. Active contractility is modelled by using a time-decreasing rest length that converges to the final value 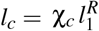 of the active branch - see Equation (2).
3. The *N*_1_ node is anchored to the substrate (*v*_1_ = 0) to allow for effective translocation of the nucleus - with speed *v*_0_.
4. No sensor *N*_2_ nodes exist in this phase.

#### 3.1 Conditions on mechanical parameters

Active contractility shall yield, in this phase, large enough values of *F*_1_ to allow for nucleus translocation, *i*.*e. F*_1_ satisfying the condition 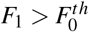. Until this condition is satisfied, the nucleus is rooted to the substrate (as the *N*_1_ node). Therefore the length of *B*_1_ remains constant - with upper bound 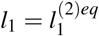 due to the elongation insured in the previous phase. The corresponding maximal contraction force in the branch is reached when fully contracted, *i*.*e*. for the minimal value 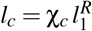 of the rest length, thus yielding the expression of the maximal contraction force

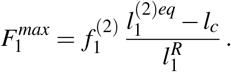

Using expression (A.3) of 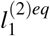, the condition 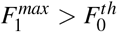 required for cell movement becomes a condition on contractility efficiency, which writes

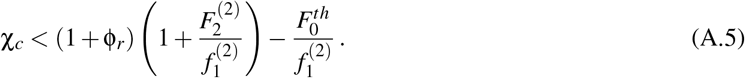

In this third and final maturation phase, cell movement based on a single *B*_1_ branch is then determined by solving the mechanical equilibrium 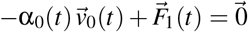 on *N*_0_ under the quasi-static approximation (see Fig. 2A) to obtain the nucleus speed :

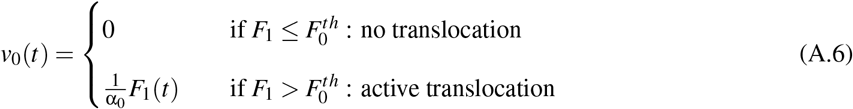

Therefore, during the contraction phase, the instantaneous speed of moving cell with a single branch lies in the interval 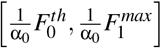, which provides a good estimate for further calibration. Remark that this estimate remains coherent for moving cell with multiple branches.

## Notes

### Competing Interest Statement

The authors have declared no competing interest.

## References

Bangasser, B. L., S. S. Rosenfeld, and D. J. Odde (2013, August). Determinants of Maximal Force Transmission in a Motor-Clutch Model of Cell Traction in a Compliant Microenvironment. Biophysical Journal 105(3), 581–592.

Bangasser, B. L., G. A. Shamsan, C. E. Chan, K. N. Opoku, E. Tüzel, B. W. Schlichtmann, J. A. Kasim, B. J. Fuller, B. R. McCullough, S. S. Rosenfeld, and D. J. Odde (2017, May). Shifting the optimal stiffness for cell migration. Nature Communications 8(1), 15313.

Bauer, A. L., T. L. Jackson, and Y. Jiang (2009, July). Topography of Extracellular Matrix Mediates Vascular Morphogenesis and Migration Speeds in Angiogenesis. PLoS Computational Biology 5(7), e1000445.

Bellomo, N., A. Bellouquid, Y. Tao, and M. Winkler (2015, August). Toward a mathematical theory of Keller–Segel models of pattern formation in biological tissues. Mathematical Models and Methods in Applied Sciences 25(09), 1663–1763.

Borau, C., R. Chisholm, P. Richmond, and D. Walker (2024). An agent-based model for cell microenvironment simulation using flamegpu2. Computers in Biology and Medicine 179, 108831.

Brückner, D. B. and C. P. Broedersz (2024, May). Learning dynamical models of single and collective cell migration: A review. Reports on Progress in Physics 87(5), 056601.

Caswell, P. T. and T. Zech (2018, October). Actin-Based Cell Protrusion in a 3D Matrix. Trends in Cell Biology 28(10), 823–834.

Chaudhuri, O., J. Cooper-White, P. A. Janmey, D. J. Mooney, and V. B. Shenoy (2020, August). Effects of extra-cellular matrix viscoelasticity on cellular behaviour. Nature 584(7822), 535–546.

Chauvière, A., I. Manifacier, C. Verdier, G. Chagnon, I. Cheddadi, N. Glade, and A. Stéphanou (2024, November). A biomechanical model for cell sensing and migration. Computer Methods in Biomechanics and Biomedical Engineering, 1–19.

Chelly, H., A. Jahangiri, M. Mireux, J. Étienne, D. K. Dysthe, C. Verdier, and P. Recho (2022, March). Cell crawling on a compliant substrate: A biphasic relation with linear friction. International Journal of Non-Linear Mechanics 139, 103897.

Chen, L., K. Painter, C. Surulescu, and A. Zhigun (2020, September). Mathematical models for cell migration: A non-local perspective. Philosophical Transactions of the Royal Society B: Biological Sciences 375(1807), 20190379.

Crossley, R. M., S. Johnson, E. Tsingos, Z. Bell, M. Berardi, M. Botticelli, Q. J. S. Braat, J. Metzcar, M. Ruscone, Y. Yin, and R. Shuttleworth (2024, March). Modeling the extracellular matrix in cell migration and morphogenesis: A guide for the curious biologist. Frontiers in Cell and Developmental Biology 12, 1354132.

De Pascalis, C. and S. Etienne-Manneville (2017, July). Single and collective cell migration: The mechanics of adhesions. Molecular Biology of the Cell 28(14), 1833–1846.

Dickinson, R. B. and R. T. Tranquillo (1993, December). Optimal estimation of cell movement indices from the statistical analysis of cell tracking data. AIChE Journal 39(12), 1995–2010.

DiMilla, P., K. Barbee, and D. Lauffenburger (1991, July). Mathematical model for the effects of adhesion and mechanics on cell migration speed. Biophysical Journal 60(1), 15–37.

Dokukina, I. V. and M. E. Gracheva (2010, June). A Model of Fibroblast Motility on Substrates with Different Rigidities. Biophysical Journal 98(12), 2794–2803.

Du Roure, O., A. Saez, A. Buguin, R. H. Austin, P. Chavrier, P. Silberzan, and B. Ladoux (2005, February). Force mapping in epithelial cell migration. Proceedings of the National Academy of Sciences 102(7), 2390–2395.

Eckert, J., Y. Abouleila, T. Schmidt, and A. Mashaghi (2021). Single Cell Micro-Pillar-Based Characterization of Endothelial and Fibroblast Cell Mechanics. Micro 2021.

Elosegui-Artola, A., X. Trepat, and P. Roca-Cusachs (2018, May). Control of Mechanotransduction by Molecular Clutch Dynamics. Trends in Cell Biology 28(5), 356–367.

Erhardt, A. H., D. Peschka, C. Dazzi, L. Schmeller, A. Petersen, S. Checa, A. Münch, and B. Wagner (2025, February). Modeling cellular self-organization in strain-stiffening hydrogels. Computational Mechanics 75(2), 875–896.

Fraley, S. I., Y. Feng, R. Krishnamurthy, D.-H. Kim, A. Celedon, G. D. Longmore, and D. Wirtz (2010, June). A distinctive role for focal adhesion proteins in three-dimensional cell motility. Nature Cell Biology 12(6), 598–604.

Ghibaudo, M., A. Saez, L. Trichet, A. Xayaphoummine, J. Browaeys, P. Silberzan, A. Buguin, and B. Ladoux (2008). Traction forces and rigidity sensing regulate cell functions. Soft Matter 4(9), 1836.

Gupton, S. L. and C. M. Waterman-Storer (2006, June). Spatiotemporal Feedback between Actomyosin and Focal-Adhesion Systems Optimizes Rapid Cell Migration. Cell 125(7), 1361–1374.

Harms, B. D., G. M. Bassi, A. R. Horwitz, and D. A. Lauffenburger (2005, February). Directional Persistence of EGF-Induced Cell Migration Is Associated with Stabilization of Lamellipodial Protrusions. Biophysical Journal 88(2), 1479–1488.

Heck, T., D. A. Vargas, B. Smeets, H. Ramon, P. Van Liedekerke, and H. Van Oosterwyck (2020, January). The role of actin protrusion dynamics in cell migration through a degradable viscoelastic extracellular matrix: Insights from a computational model. PLOS Computational Biology 16(1), e1007250.

Kaverina, I., O. Krylyshkina, and J. Small (2002, July). Regulation of substrate adhesion dynamics during cell motility. The International Journal of Biochemistry & Cell Biology 34(7), 746–761.

Keijzer, K. A. E., E. Tsingos, and R. M. H. Merks (2025, January). How cells align to structured collagen fibrils: A hybrid cellular Potts and molecular dynamics model with dynamic mechanosensitive focal adhesions. Frontiers in Cell and Developmental Biology 12, 1462277.

Kicheva, A., P. Pantazis, T. Bollenbach, Y. Kalaidzidis, T. Bittig, F. Jülicher, and M. González-Gaitán (2007, January). Kinetics of Morphogen Gradient Formation. Science 315(5811), 521–525.

Larsen, M., V. V. Artym, J. A. Green, and K. M. Yamada (2006, October). The matrix reorganized: Extracellular matrix remodeling and integrin signaling. Current Opinion in Cell Biology 18(5), 463–471.

Liang, H., W. Fang, and X.-Q. Feng (2024). A multiscale dynamic model of cell–substrate interfaces. Journal of the Mechanics and Physics of Solids 189, 105725.

Marchello, R., A. Colombi, L. Preziosi, and C. Giverso (2024, February). A non local model for cell migration in response to mechanical stimuli. Mathematical Biosciences 368, 109124.

Mech, D. J. and M. S. Rizvi (2024, March). In-silico modeling of the micromechanics of fibrous scaffolds and stiffness sensing by cells. Biomedical Materials 19(2), 025035.

Merino-Casallo, F., M. J. Gomez-Benito, R. Martinez-Cantin, and J. M. Garcia-Aznar (2022, June). A mechanistic protrusive-based model for 3D cell migration. European Journal of Cell Biology 101(3), 151255.

Metzcar, J., B. S. Duggan, B. Fischer, M. Murphy, R. Heiland, and P. Macklin (2025). A simple framework for agent-based modeling with extracellular matrix. Bulletin of Mathematical Biology 87(3), 43.

Noël, V., M. Ruscone, R. Shuttleworth, and C. K. Macnamara (2024, October). PhysiMeSS - a new physiCell addon for extracellular matrix modelling. Gigabyte 2024, gigabyte135.

Palecek, S. P., J. C. Loftus, M. H. Ginsberg, D. A. Lauffenburger, and A. F. Horwitz (1997). Integrin-ligand binding properties govern cell migration speed through cell-substratum adhesiveness. Nature 385(6616), 537–540.

Pawluchin, A. and M. Galic (2022, December). Moving through a changing world: Single cell migration in 2D vs. 3D. Frontiers in Cell and Developmental Biology 10, 1080995.

Peschetola, V. (2011, November). Détermination des forces de traction au cours de la migration de cellules cancéreuses sur des gels. Ph. D. thesis, Université Grenoble-Alpes.

Pham, J. T., L. Xue, A. Del Campo, and M. Salierno (2016, July). Guiding cell migration with microscale stiffness patterns and undulated surfaces. Acta Biomaterialia 38, 106–115.

Reinhardt, J. W., D. A. Krakauer, and K. J. Gooch (2013, July). Complex Matrix Remodeling and Durotaxis Can Emerge From Simple Rules for Cell-Matrix Interaction in Agent-Based Models. Journal of Biomechanical Engineering 135(7), 071003.

Rens, E. G. and R. M. Merks (2020, September). Cell Shape and Durotaxis Explained from Cell-Extracellular Matrix Forces and Focal Adhesion Dynamics. iScience 23(9), 101488.

Roux, C., A. Duperray, V. M. Laurent, R. Michel, V. Peschetola, C. Verdier, and J. Étienne (2016, October). Prediction of traction forces of motile cells. Interface Focus 6(5), 20160042.

Schlüter, D. K., I. Ramis-Conde, and M. A. Chaplain (2012, September). Computational Modeling of Single-Cell Migration: The Leading Role of Extracellular Matrix Fibers. Biophysical Journal 103(6), 1141–1151.

Scianna, M., L. Preziosi, and K. Wolf (2013). A Cellular Potts model simulating cell migration on and in matrix environments. Mathematical Biosciences and Engineering 10(1), 235–261.

Shellard, A. and R. Mayor (2021, December). Collective durotaxis along a self-generated stiffness gradient in vivo. Nature 600(7890), 690–694.

Shemesh, T., A. D. Bershadsky, and M. M. Kozlov (2012, April). Physical Model for Self-Organization of Actin Cytoskeleton and Adhesion Complexes at the Cell Front. Biophysical Journal 102(8), 1746–1756.

Small, J. V. and G. P. Resch (2005, October). The comings and goings of actin: Coupling protrusion and retraction in cell motility. Current Opinion in Cell Biology 17(5), 517–523.

Sopher, R. S., H. Tokash, S. Natan, M. Sharabi, O. Shelah, O. Tchaicheeyan, and A. Lesman (2018, October). Nonlinear Elasticity of the ECM Fibers Facilitates Efficient Intercellular Communication. Biophysical Journal 115(7), 1357–1370.

Stéphanou, A., S. L. Floc’h, and A. Chauvière (2015). Hybrid modelling of mechanical cues in cell migration. ITM Web of Conferences 5, 00012.

Stéphanou, A., S. McDougall, A. Anderson, and M. Chaplain (2005, May). Mathematical modelling of flow in 2D and 3D vascular networks: Applications to anti-angiogenic and chemotherapeutic drug strategies. Mathematical and Computer Modelling 41(10), 1137–1156.

Stéphanou, A., E. Mylona, M. Chaplain, and P. Tracqui (2008, August). A computational model of cell migration coupling the growth of focal adhesions with oscillatory cell protrusions. Journal of Theoretical Biology 253(4), 701–716.

Svitkina, T. (2018, January). The Actin Cytoskeleton and Actin-Based Motility. Cold Spring Harbor Perspectives in Biology 10(1), a018267.

Verdier, C., J. Etienne, A. Duperray, and L. Preziosi (2009, November). Review: Rheological properties of biological materials. Comptes Rendus. Physique 10(8), 790–811.

Voss-Böhme, A. (2012, September). Multi-Scale Modeling in Morphogenesis: A Critical Analysis of the Cellular Potts Model. PLoS ONE 7(9), e42852.

Wössner, V., O. M. Drozdowski, F. Ziebert, and U. S. Schwarz (2024, July). Active gel model for one-dimensional cell migration coupling actin flow and adhesion dynamics. New Journal of Physics 26(7), 073039.

Yamada, K. M. and M. Sixt (2019, December). Mechanisms of 3D cell migration. Nature Reviews Molecular Cell Biology 20(12), 738–752.

Zaidel-Bar, R., M. Cohen, L. Addadi, and B. Geiger (2004, June). Hierarchical assembly of cell–matrix adhesion complexes. Biochemical Society Transactions 32(3), 416–420.

Zhu, H., R. Miao, J. Wang, and M. Lin (2024, March). Advances in modeling cellular mechanical perceptions and responses via the membrane-cytoskeleton-nucleus machinery. Mechanobiology in Medicine 2(1), 100040.

